# Stable isotope tracing in human plasma-like medium reveals metabolic and immune modulation of the glioblastoma microenvironment

**DOI:** 10.1101/2023.05.29.542774

**Authors:** Milan R. Savani, Mohamad El Shami, Lauren C. Gattie, Bailey C. Smith, William H. Hicks, Skyler S. Oken, Lauren G. Zacharias, Misty S. Martin-Sandoval, Eric Y. Montgomery, Yi Xiao, Diana D. Shi, Jeremy N. Rich, Timothy E. Richardson, Pascal O. Zinn, Bradley C. Lega, Thomas P. Mathews, Ralph J. DeBerardinis, Samuel K. McBrayer, Kalil G. Abdullah

## Abstract

**Background:** In vivo stable isotope tracing is useful for natively surveying glioma metabolism but can be difficult to implement. Stable isotope tracing is tractable using in vitro glioma models, but most models lack nutrient conditions and cell populations relevant to human gliomas. This limits our ability to study glioma metabolism in the presence of an intact tumor microenvironment (TME) and immune-metabolic crosstalk.

**Methods:** We optimized an in vitro stable isotope tracing approach for human glioma explants and glioma stem-like cell (GSC) lines that integrates human plasma-like medium (HPLM). We performed ^15^N_2_-glutamine tracing in GSC monocultures and human IDH-wildtype glioblastoma explants and developed an analytical framework to evaluate microenvironment-dependent metabolic features that distinguish them. We also conducted spatial transcriptomics to assess transcriptional correlates to metabolic activities.

**Results:** HPLM culture preserved glioma explant viability and stemness while unmasking metabolic and immune programs suppressed by conventional culture conditions. Stable isotope tracing in HPLM revealed TME-dependent and TME-independent features of tumor metabolism. Tissue explants recapitulated tumor cell-intrinsic metabolic activities, such as synthesis of immunomodulatory purines. Unlike GSC monocultures, tissue explants captured tumor cell-extrinsic activities associated with stromal cell metabolism, as exemplified by astrocytic GDP-mannose production in heterocellular explants. Finally, glioma explants displayed tumor subtype-specific metabolic reprogramming, including robust pyrimidine degradation in mesenchymal cells.

**Conclusions:** We present a tractable approach to assess glioma metabolism in vitro under physiological nutrient levels and in the presence of an intact TME. This platform opens new avenues to interrogate glioma metabolism and its interplay with the immune microenvironment.

**Importance of the Study:** Metabolic reprogramming is a hallmark of tumor biology, but in vitro studies of glioma metabolism often fail to replicate the nutrient complexity and cellular heterogeneity of the TME. We developed a method to perform stable isotope tracing in glioma explants grown in HPLM to analyze metabolism in a nutrient context that reflects in vivo conditions. By comparing metabolic activities between glioma cell monocultures and explanted tumor tissues, our approach captures features of tumor metabolism that are driven by the microenvironment. We show that HPLM not only sustains cell fitness and identity in tumor explants but also evokes distinct metabolic patterns and immune activation signatures repressed by standard culture conditions. Our approach offers a tractable and scalable way to study tumor cell intrinsic and microenvironmental metabolism in faithful tissue culture glioma models, complementing powerful yet low-throughput in vivo stable isotope tracing approaches.

**Key Points:**

- HPLM supports culture of glioma explants and stimulates metabolic and immune transcriptional responses.
- Stable isotope tracing in glioma explants reveals contributions of tumor cells, stromal cells, and gene expression programs to tumor metabolism.

## INTRODUCTION

Metabolic reprogramming is a hallmark of cancer biology that provides energy and substrates for cell growth^1,2^. There has been substantial interest in the metabolic adaptations that enable glioma cells to grow and proliferate, which has resulted in the nomination of new therapeutic targets^3–11,11–17^. Tumor-relevant biochemical networks are flexible and sensitive to concentrations of environmental nutrients and the metabolic activities of neighboring cells^18,19^. The gold standard for investigating glioma metabolism while capturing microenvironmental interactions is to utilize in vivo stable isotope tracing in mouse brain tumor models or in patients^20–26^. These experiments are technically challenging, costly, and time-consuming.

However, performing stable isotope tracing in more accessible systems, such as cell culture models, may not capture vital characteristics of tumor metabolism, including heterocellular interactions of the tumor microenvironment (TME) and physiologic nutrient levels.

Culture media that more accurately reflect physiologic nutrient availability—such as human plasma-like medium (HPLM)—have recently been applied to cancer models and reveal marked effects on cellular metabolism and nutrient utilization^3,27–29^. HPLM contains human-relevant concentrations of typical media components such as glucose, amino acids, and salt ions, as well as some metabolites absent from commonly used culture media. Mimicking physiological nutrient conditions has profound effects on cellular metabolism, and analysis of tumor metabolism with stable isotope-labeled metabolites in HPLM can reveal metabolic activities that may be obscured by standard cell culture media^3,27^. However, efforts to transition in vitro glioma models from standard to physiological media remain limited, potentially due to concerns that altered nutrient conditions may induce GSC differentiation or death. Therefore, it is not clear whether HPLM culture can be exploited to unmask interactions between metabolism and the immune microenvironment in glioma, as has been demonstrated in other contexts^28^.

One promising system for modeling glioma metabolism in vitro has been the use of surgically explanted glioma tissue, particularly Surgically eXplanted Organoids (SXOs). SXOs are efficiently created without single-cell dissociation of the resected tumor tissue, allowing maintenance of local cytoarchitecture^30^. These models recapitulate parental tumor features such as cellular heterogeneity, cell-cell and cell-stroma interactions, gene expression and mutational profiles, and treatment response^30–32^. While SXOs offer advantages over conventional models, they are typically cultured in non-physiologic media such as Glioma Organoid Complete (GOC) medium, likely altering nutrient-sensitive metabolic programs. Integrating HPLM culture with glioma organoid modeling efforts may improve fidelity of metabolism studies while retaining the structural complexity of the native tumor microenvironment.

Here, we integrate an adapted formulation of HPLM with stable isotope tracing in both GSC monocultures and IDH-wildtype GBM tissue explants. We develop a quantitative labeling score to compare metabolic pathway activity between models and pair these data with spatial transcriptomics to identify nutrient-sensitive transcriptional programs. This approach establishes a physiologic in vitro model of glioma metabolism that complements existing strategies to study tumor cell-intrinsic and microenvironmental metabolic activities.

## MATERIALS AND METHODS

### Human Subjects

This study was conducted according to the principles of the Declaration of Helsinki. Patient tissue and blood were collected following ethical and technical guidelines on the use of human samples for biomedical research at University of Texas Southwestern Medical Center after informed patient consent under protocols approved by the University of Texas Southwestern Medical Center’s IRB (STU 022011-070 and STU 092014-026).

### Primary Cell Culture

BT054 (female, RRID: CVCL_N707) cells were obtained from S. Weiss at the University of Calgary^33^. TS516 (sex unknown, RRID: CVCL_A5HY) and TS603 (sex unknown, RRID: CVCL_A5HW) cells were obtained from I. Mellinghoff at Memorial Sloan-Kettering Cancer Center^34^. HK157 (female), HK211 (female), HK213 (male), HK252 (male), and HK308 (male) cells were obtained from H. Kornblum at the University of California Los Angeles^35^. MGG152 (male) cells were obtained from D. Cahill at Massachusetts General Hospital^36^. BT054, HK157, TS516, TS603, UTSW5, UTSW63, and UTSW71 cells were cultured in NeuroCult NS-A Basal Medium (Human) with Proliferation Supplement (StemCell Technologies 05751), supplemented with EGF (20 ng/mL, GoldBio 1150-04-100), bFGF (20 ng/mL, GoldBio 1140-02-10), heparin (2 µg/mL, STEMCELL Technologies 07980), penicillin/streptomycin (100 U/mL and 100 μg/mL, respectively, Gibco 15140148), amphotericin B (250 ng/mL, Gemini Bio-Products 400104), and Plasmocin (250 ng/mL, Invitrogen ant-mpp). HK157 (when used for RNA sequencing assays), HK211, HK252, and HK308 cells were cultured in DMEM-F12 medium (Gibco 11320033) supplemented with glutamine (3mM, Gibco 25030081), B27 (1×, Gibco 17504044), EGF (20 ng/mL, GoldBio 1150-04-100), bFGF (20 ng/mL, GoldBio 1140-02-10), heparin (2 µg/mL, STEMCELL Technologies 07980), penicillin/streptomycin (50 U/mL and 50 μg/mL, respectively, Gibco 15140148), amphotericin B (125 ng/mL, Gemini Bio-Products 400104), and Plasmocin (250 ng/mL, Invitrogen ant-mpp).

MGG152 cells were cultured in Neurobasal Medium (Gibco 21103049) supplemented with glutamine (3mM, Gibco 25030081), B27 (1×, Gibco 17504044), N2 (0.25×, Gibco 17502048), EGF (20 ng/mL, GoldBio 1150-04-100), bFGF (20 ng/mL, GoldBio 1140-02-10), heparin (2 µg/mL, STEMCELL Technologies 07980), penicillin/streptomycin (50 U/mL and 50 μg/mL, respectively, Gibco 15-140-148), amphotericin B (125 ng/mL, Gemini Bio-Products 400104), and Plasmocin (0.25 µg/mL, Invitrogen ant-mpp). GSCs were cultured in 5% CO2 and at ambient oxygen at 37°C. All GSCs were routinely evaluated for mycoplasma contamination and confirmed to be negative.

### Explant SXO Creation and Culture in GOC

Tumor tissue was collected from the operating room, directly suspended in ice cold Hibernate A (BrainBits HA), and transferred to the laboratory on ice within 30 minutes of explantation. Tumor pieces were moved into RBC lysis buffer (Thermo Fisher 00433357) and incubated at room temperature for 10 minutes with rocking. Tumor pieces were then washed with Hibernate A containing Glutamax (2 mM, Thermo Fisher 35050061), penicillin/streptomycin (100 U/mL and 100 μg/mL, respectively, Gibco 15140148), and Amphotericin B (250 ng/mL, Gemini Bio-Products 400104). Tissues were cut into SXOs using a 750 µm^2^ internal diameter needle (SAI Infusion Technologies B18-150) and plated, one per well, in a 24-well ultra-low adherence plate (Corning 3473) in 1 mL Glioma Organoid Complete Medium (GOC)^30^. Stocks of GOC were used within a maximum of 1 week after preparation. Plates were rotated at 120 rpm in a humidified incubator at 37°C, 5% CO_2_, and 21% oxygen. GOC was replaced in SXO cultures every 48 hours. All SXOs were cultured for a minimum of eight weeks prior to experimentation.

### Culture in Glioma Stem-Like Cell (GSC) HPLM

HPLM was prepared as previously described^27,28,37^ but without the addition of dialyzed fetal bovine serum and glutamate. HPLM was then supplemented with B27 (1×, Gibco 17504044), N2 (0.25×, Gibco 17502048), EGF (20 ng/mL, GoldBio 1150-04-100), bFGF (20 ng/mL, GoldBio 1140-02-10), heparin (2 µg/mL, STEMCELL Technologies 07980), penicillin/streptomycin (100 U/mL and 100 μg/mL, respectively, Gibco 15-140-148), amphotericin B (250 ng/mL, Gemini Bio-Products 400104), and Plasmocin (250 ng/mL, Invitrogen ant-mpp). For isotopically labeled GSC HPLM, GSC HPLM was prepared as above, substituting equimolar either ^15^N_2_ glutamine (Cambridge Isotope Laboratories NLM-1328) or amide-^15^N glutamine (Cambridge Isotope Laboratories NLM-557) for unlabeled glutamine. Stocks of GSC HPLM were used within a maximum of 1 week after preparation. Prior to experiments conducted in GSC HPLM, GSCs were first cultured for 24 hours in a mixture of 50% NeuroCult and 50% GSC HPLM, then for 24 hours in 100% GSC HPLM.

### Explant Culture in SXO HPLM

HPLM was prepared as previously described^27,28,37^ but without addition of dialyzed fetal bovine serum. HPLM was then supplemented with B27 without Vitamin A (1×, Gibco 12587010), N2 (1×, Gibco 17502048), 2-mercaptoethanol (55µM, Thermo Fisher BP176-100), human insulin (2.375-2.875µg/mL, Sigma Aldrich I9278). For isotopically labeled SXO HPLM, SXO HPLM was prepared as above, substituting equimolar ^15^N_2_ glutamine (Cambridge Isotope Laboratories NLM-1328) for unlabeled glutamine. Stocks of SXO HPLM were used within a maximum of 1 week after preparation. Prior to experiments conducted in SXO HPLM, SXOs were first cultured for 24 hours in a mixture of 50% GOC and 50% SXO HPLM, then for 24 or 120 hours in 100% SXO HPLM, replacing HPLM every 24 hours.

### Histology and Immunohistochemistry

SXOs were fixed in 10% formalin for 1 hour, washed, and suspended in 70% ethanol. Samples were embedded in paraffin and sectioned at 4μm prior to staining. Embedding, sectioning, histology, immunohistochemistry, and digital image acquisition were performed by HistoWiz, Inc. (histowiz.com). Images were processed using FIJI (1.53f51, imagej.net/software/fiji, RRID:SCR_002285). Nuclei and diaminobenzidine positivity in immunohistochemical stains were quantified with a semi-automated trained object classifier, implemented in QuPath (0.3.1, qupath.github.io, RRID:SCR_018257).

### Liquid Chromatography-Mass Spectrometry (LC-MS)

For stable isotope tracing experiments, SXOs were washed in ice cold saline prepared in LC-MS grade water (Fisher Scientific W6500), then snap-frozen and stored at -80°C until analysis. Accurate masses were obtained using an analytical balance. 80% LC-MS grade acetonitrile (Fisher Scientific A9554) prepared in LC-MS grade water was added at 100µL per mg of tissue to snap-frozen SXOs on ice, followed by homogenization by manual agitation. Homogenate was vortexed for 20 minutes at 4°C, then centrifuged for 10 minutes at 21,100 × g at 4°C. Supernatant was transferred to a fresh microcentrifuge tube and centrifuged again for 10 minutes at 21,100 × g at 4°C. For GSCs, neurospheres were harvested from 6-well plates, followed by the addition of ice-cold saline prepared in LC-MS grade water. GSCs were transferred to microcentrifuge tubes and centrifuged for 1 minute at 21,100 × g at 4°C. Supernatant was aspirated, then cell pellets were snap-frozen and stored at -80°C. Metabolites were extracted in 80% Acetonitrile at a concentration of 1,000 cells/µL, vortexed for 20 minutes at 4°C, and centrifuged for 10 minutes at 21,100 × g at 4°C. Supernatant was transferred to a fresh microcentrifuge tube and centrifuged again for 10 minutes at 21,100 × g at 4°C. For both SXOs and GSCs, final supernatant was transferred to a glass vial for LC-MS analysis. Data acquisition of isotopically-labeled metabolites was performed as described previously^38,39^. Briefly, 10µL of SXO sample or 20uL of GSC sample was injected and analyzed with an Q Exactive™ HF-X orbitrap mass spectrometer (Thermo Fisher) coupled to a Vanquish ultra-high performance liquid chromatography system (Thermo Fisher). Peaks were integrated using El-Maven software (0.12.0, Elucidata). Total ion counts were quantified using TraceFinder software (5.1 SP2, Thermo Fisher). Peaks were normalized to total ion counts using the R statistical programming language. Correction for natural abundance of ^15^N isotopes was accomplished using the R script Accucor^40^. Total fractional enrichment was calculated by subtracting the fractional abundance of the M+0 isotopologue from 1 for each metabolite.

### Derivation of Metabolite Labeling Score

Total fractional enrichment of all quantified metabolites was calculated as described above. Metabolites were filtered for total labeling > 0 and adequate total pool size. Each metabolite’s total labeling was divided by label accumulation in glutamate to normalize and derive the Metabolite Labeling Score.

### Metabolite Set Enrichment Analysis

Quantitative enrichment analysis was performed using MetaboAnalyst (6.0)^41,42^. Metabolite Labeling Scores were compared between SXO210 and, separately, UTSW63, TS516, and HK157 without normalization, filtering, or scaling.

### Digital Spatial Profiling (DSP) Assay

Spatial expression profiles were analyzed using GeoMx DSP (NanoString Technologies, RRID:SCR_021660). After deparaffinization and rehydration, 4μm formalin-fixed paraffin-embedded (FFPE) tissue slides were hybridized and incubated at 37°C overnight with morphological immunofluorescent biomarkers and DSP probes from the Cancer Transcriptome Atlas panel (NanoString Technologies). SYTO-13, GFAP, and CD45 were used as visualization markers. Slides were scanned on the GeoMx DSP instrument to produce a digital image displaying fluorescent visualization markers. Spatially resolved ROIs were selected based on fluorescent markers. DSP probes conjugated to target-specific activated oligos were collected for each ROI and aliquoted into 96-well plates. Collection plates were dehydrated at 65°C for 1-2 hours on a thermal cycler with an open top and a breathable AeraSeal film (Excel Scientific, A9224). Samples were reconstituted with diethyl pyrocarbonate-treated RNase/DNase-free water, and library preparation was completed with eighteen amplification cycles. Libraries were quantified on an Agilent 4200 TapeStation and pooled for sequencing on an Illumina NextSeq 2000 with a P3 50 flow cell. Whole transcriptome gene sequencing was performed on individual ROI tubes by the University of Pittsburgh Health Sciences Sequencing Core, Rangos Research Center, UPMC Children’s Hospital of Pittsburgh. Resulting FASTQ files were decoded and processed into count files using the NanoString GeoMx NGS Pipeline (2.0.21, NanoString Technologies) in the Illumina BaseSpace Sequencing Hub. Count files were uploaded to the GeoMx DSP instrument and indexed to corresponding slide scans for analysis. Gene counts were mapped to selected ROIs and normalized after a quality check for the following parameters in each ROI: raw read count > 1,000, >80% of reads aligned, sequencing saturation >50%, negative probe geometric mean >10 in background, count of nuclei per ROI >200, and surface area per ROI <16,000 µm^2^. CIBERSORT (RRID:SCR_016955), a deconvolution algorithm built on nine normalized gene expression profiles to characterize cell composition^43^, was used to estimate cell population proportions based on the leukocyte signature matrix 22. CIBERSORT was run for 1,000 permutations, and samples with a CIBERSORT *p* value below 0.05 were included for subsequent analyses. Gene expression analysis was performed using GeoMx Analysis suite (2.4.2.2, RRID:SCR_023424). Gene set enrichment analysis (GSEA; 4.2.3; Broad Institute, Inc., Massachusetts Institute of Technology, and Regents of the University of California; RRID:SCR_003199) based the Molecular Signatures Database (RRID:SCR_016863) was performed to compare SXOs cultured in HPLM to SXOs cultured in GOC. Clinical next-generation sequencing of the parental tumor was performed on FFPE-preserved tissue and paired germline DNA from patient saliva by the UT Southwestern Clinical NGS laboratory (CLIA ID 45D0861764).

### Drug Treatment

Gimeracil (Cayman Chemical Company 16525) was used at 30 µM and TNFα (Sigma-Aldrich H8916) was used at 10 ng/mL where indicated.

### CD44 Quantification

Cells were stained with CD44-FITC (Miltenyi 130-113-341) and CD24- APC (Miltenyi 130-095-954) or the isotype controls IgG1-FITC (Miltenyi 130-113-437) and IgG1-APC (Miltenyi 130-113-196) according to manufacturer’s instructions. Cells were analyzed using an LSR Fortessa (BD Biosciences) flow cytometer.

### TCGA Expression Analysis

The TCGA Lower Grade Glioma (LGG)^44^ and GBM^45^ datasets were used to assess correlation between *CD44* and *DPYD* or *HPRT1* expression levels in human gliomas (https://www.cbioportal.org).

### Quantification and Statistical Analysis

SXOs were allocated to experiments randomly and samples were processed in an arbitrary order. All statistical tests were two-sided, where applicable. Student’s *t*-test was used to assess the statistical significance of a difference between the two groups. Linear regression analysis and Pearson’s correlation coefficient were used to assess correlation between the two groups. Statistical analyses were performed with GraphPad Prism (10.4.1, GraphPad Software, LLC, RRID:SCR_002798) and included both descriptive statistics as well as tests of statistical significance. All data are plotted as mean ± standard error of the mean. For all tests, *p-*values less than 0.05 were considered statistically significant.

## RESULTS

### HPLM preserves glioma cytoarchitecture and reveals nutrient-responsive transcriptional programs

We developed a comprehensive experimental approach to assess the feasibility and utility of applying HPLM culture and SXO glioma models to study glioma metabolism (Figure 1). Our approach included comparisons of explants cultured in conventional GOC medium with those grown in HPLM. We conducted histological, metabolic, and spatial transcriptomic analyses of glioma SXOs and benchmarked our findings against GSC monocultures to discriminate tumor cell-intrinsic and -extrinsic properties reflected in explant models. To do so, we established SXOs from surgically resected IDH-wildtype glioblastoma (GBM) tissue (Table S1).

**Figure 1.**
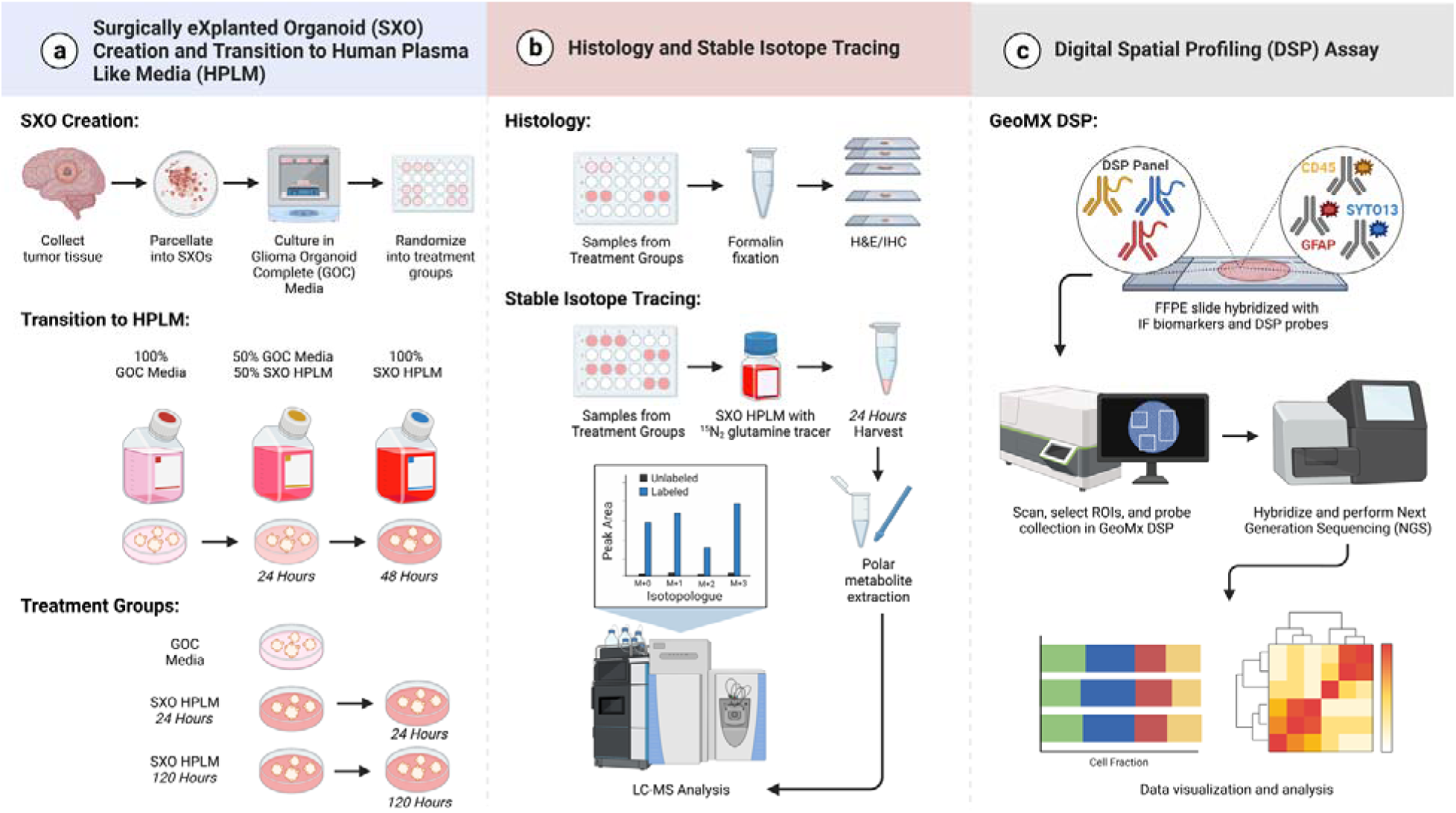
Experimental overview. **(a)** Glioblastoma Surgically eXplanted Organoid (SXO) creation and transition to Human Plasma-Like Medium (HPLM). Human brain tumor tissue was collected directly from the operating room, manually parcellated, and cultured as explants. SXOs were cultured in Glioma Organoid Complete Medium with glutamate (GOC), then randomized to culture in GOC or SXO HPLM. For HPLM culture, SXOs were first transferred into media containing 50% GOC and 50% SXO HPLM for 24 hours, then preconditioned in 100% SXO HPLM for either an additional 24 or 120 hours. **(b)** Histology and stable isotope tracing. *Histology:* SXOs were fixed in neutral buffered formalin, then were stained for hematoxylin and eosin (H&E), Ki67, and Sox2. *Stable Isotope Tracing:* SXOs cultured in SXO HPLM were transferred into SXO HPLM containing ^15^N_2_-glutamine. After 24 hours, polar metabolites were harvested and subjected to liquid chromatography-mass spectrometry (LC-MS). **(c)** NanoString Digital Spatial Profiling (DSP) Assay. Formalin-fixed paraffin-embedded tissue slides were deparaffinized and stained with fluorescent antibodies (SYTO-13 to identify DNA, GFAP to identify tumor cells, and CD45 to identify immune cells). Spatially resolved regions of interest (ROIs) were selected for analysis based on fluorescent images. Oligo-tagged probes for each ROI were used for spatial transcriptomics analysis. Figure created with BioRender.

We first compared the viability and cytoarchitecture of glioma explants cultured in an SXO- adapted formulation of HPLM to those cultured in conventional GOC medium (Figures 2A-F). Histological evaluation by a board-certified neuropathologist (TER) revealed that explants maintained hallmark features of GBM—including necrosis, microvascular proliferation, high mitotic index, and cellular atypia—regardless of culture condition. Quantitative analysis of nuclei demonstrated that SXOs cultured in SXO HPLM for both 24 and 120 hours maintained similar cell density to those cultured in GOC (Figure 2G). These results suggest that explants do not acutely lose integrity or cellularity upon transition to HPLM culture. Using Ki67 as a marker of cellular proliferation^46^ (Figures 2H-M) we found that the SXOs cultured in GOC and SXO HPLM for 24 hours exhibited similar proliferation, while the SXOs cultured in SXO HPLM for 120 hours exhibited a slight proliferation decrease (Figure 2N). We also evaluated explant tissues for expression of the stemness marker Sox2^47^ (Figures 2O-T). Sox2-positive cell frequencies were similar between all groups (Figure 2U), suggesting that HPLM culture does not cause GSC differentiation over durations spanning one to five days.

**Figure 2.**
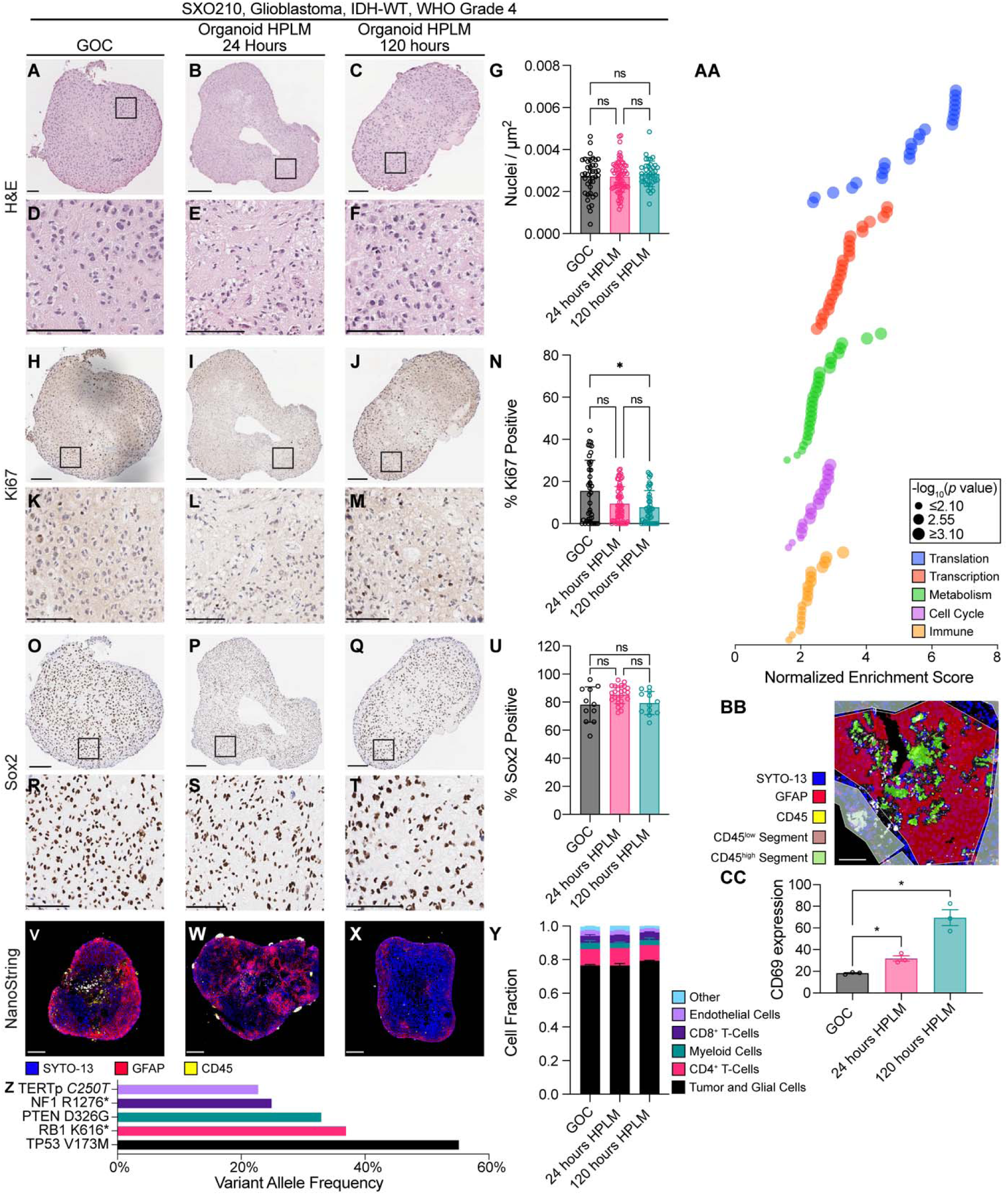
SXO histology and transcriptional response after culture in SXO HPLM. (A-F) H&E of representative SXOs cultured in **(A)** GOC, **(B)** SXO HPLM for 24 hours, and **(C)** SXO HPLM for 120 hours, which are shown at high magnification in **(D-F)**. Scale bars = 100 μm. **(G)** Imaging-based quantification of cell density by evaluation of nuclei/µm^2^. ns = not significant (Tukey’s multiple comparisons test). **(H-M)** Ki67 immunohistochemistry (IHC) of representative SXOs cultured in **(H)** GOC, **(I)** SXO HPLM for 24 hours, and **(J)** SXO HPLM for 120 hours, which are shown at high magnification in **(K-M)**. Scale bars = 100 μm. **(N)** Quantification of Ki67-positive cells. ns = not significant, \**p* < 0.05 (Dunnett’s T3 multiple comparisons test). **(O- T)** Sox2 IHC of representative SXOs cultured in **(O)** GOC, **(P)** SXO HPLM for 24 hours, and **(Q)** SXO HPLM for 120 hours, which are shown at high magnification in **(R-T)**. Scale bars = 100 μm. **(U)** Quantification of Sox2-positive cells. ns = not significant (Dunnett’s T3 multiple comparisons test). **(V-X)** NanoString morphology scans demonstrating immunofluorescent staining for SYTO- 13 (nuclei, blue), GFAP (tumor and glial cells, red), and CD45 (immune cells, yellow) in representative SXOs cultured in **(V)** GOC, **(W)** SXO HPLM for 24 hours, and **(X)** SXO HPLM for 120 hours. Scale bars = 100 μm. **(Y)** CIBERSORT imputed cell fractions from NanoString gene expression data. **(Z)** Variant allele frequencies for identified variants of strong clinical significance and variants of possible clinical significance by a clinical next-generation sequencing assay using DNA and RNA sequencing of FFPE of parental tumor for SXO210 compared to patient germline DNA from saliva. **(AA)** Gene set enrichment analysis (GSEA) of geometric mean normalized NanoString sequencing data from SXO210 cultured in SXO HPLM for 120 hours versus SXO210 cultured in GOC, grouped by pathway category. *p* values are determined by GSEA. **(BB)** NanoString morphology scan in a representative immune- segmented SXO210 ROI, with highlights demonstrating segmentation (CD45^low^ red, CD45^high^ green). Scale bar = 100 μm. **(CC)** Quantification of relative CD69 expression in CD45+ segments of SXO210 ROIs. Expression data are geometric mean normalized and *z-*scored relative to other genes. ns = not significant, \**p* < 0.05 (unpaired *t*-test). Data are presented as means ± standard error of the mean.

To evaluate cellular heterogeneity and gene expression programs, we performed NanoString-based spatial transcriptomics using immunofluorescent markers SYTO-13 (to mark DNA)^48^, GFAP (to mark astrocytes and malignant cells)^49^, and CD45 (to mark immune populations)^50^ of SXOs cultured in GOC or SXO HPLM for 24 or 120 hours (Figures 2V-X). Morphology scans for NanoString analysis were collected and used to select regions of interest (ROIs) containing either diverse heterocellular populations or predominantly immune cells for spatially resolved sequencing. Using sequencing results from heterocellular SXO ROIs, we employed the CIBERSORT algorithm^43^ to impute cell identities and estimate fractions of cell types present in each sample (Figure 2Y). SXOs contained a diverse, spatially variable population of constituent cells, consistent with prior work from our and other groups^30,32^. Cellular composition of SXOs did not vary between culture conditions. Clinical next-generation sequencing of the parental tumor used to generate SXOs revealed mutations of likely clinical significance affecting *TP53*, *RB1*, *PTEN*, *NF1*, and *TERT* genes (Figure 2Z). Using NanoString sequencing results, we performed gene set enrichment analysis. Categories of enriched genes in 120 hours of HPLM culture included those involved in translation, gene transcription, cellular metabolism, the cell cycle, and immunologic pathways (Figure 2AA and Table S2). Prior reports have indicated that culture in HPLM increases markers of T cell activation compared to traditional cell culture.^28^ We used sequencing results from immune-predominant ROIs (Figure 2BB) to investigate expression of CD69, a marker of activation of several T-cell populations^51^. CD69 expression increased in a time-dependent manner following transition to HPLM culture (Figure 2CC). Together, these data indicate that SXOs cultured in HPLM maintain viability, recapitulate the cellular heterogeneity of parental tumors, engage transcriptional programs that are dampened in conventional tissue culture media, and display heightened immune cell activity.

### Stable isotope tracing in HPLM to compare metabolic patterns in GSC monocultures and glioma explants

We performed ^15^N_2_-glutamine stable isotope tracing in explants cultured in SXO HPLM. SXO210 organoids were preconditioned in tracer-free HPLM for either 24 or 120 hours and then transferred to tracer-free or ^15^N_2_-glutamine-containing HPLM for 24 hours. We employed LC-MS to quantify ^15^N-labeled and unlabeled metabolites in extracts from these organoids. First, we confirmed intra-organoid accumulation of isotopic label in glutamine (Figure 3A). We compared label accumulation from ^15^N_2_ glutamine in all metabolites with detectable labeling between SXOs preconditioned for 24 and 120 hours in HPLM. Importantly, we observed minimal metabolite labeling in explants cultured in tracer-free HPLM (Figures 3A, 3C). Linear regression analysis revealed no difference in total labeling, with an *r*^2^ of 0.9427 indicating strong concordance between conditions (Figure 3B). In selected metabolites representing a diverse array of biosynthetic pathways (including nonessential amino acid biosynthesis, branched-chain amino acid transamination, redox homeostasis, nucleotide synthesis, and the urea cycle), there were no differences in labeling between 24- and 120-hour preconditioned samples (Figure 3C). Therefore, we selected the 120-hour HPLM preconditioning approach for further analysis.

**Figure 3.**
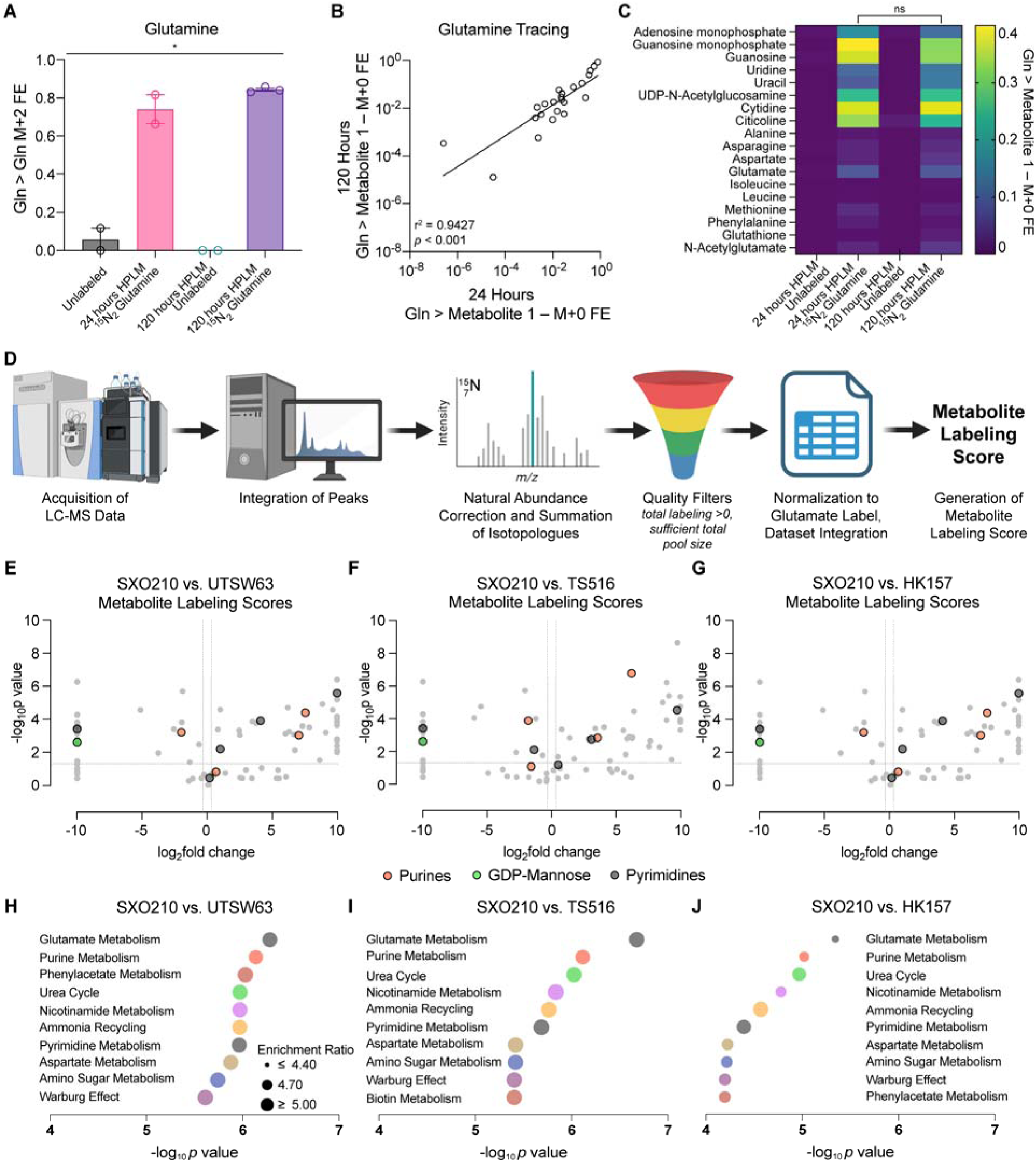
Stable isotope tracing in SXOs and GSCs reveals tumor cell-intrinsic and - extrinsic features of glioma metabolism. (A-C) ^15^N_2_-glutamine (Gln) stable isotope tracing assay in SXO210. *n* = 2 for unlabeled and 24-hour HPLM preconditioning groups, *n* = 3 for 120- hour HPLM preconditioning group. **(A)** Fractional enrichment (FE) of the M+2 isotopologue of glutamine in glutamine pool. \**p* < 0.05 (one-way ANOVA). **(B)** Correlation between label accumulation in all detected and labeled metabolites between 24-hour and 120-hour HPLM preconditioning. *r* = Pearson’s correlation coefficient. *p* value was determined by simple linear regression analysis**. (C)** Total FE (1 – M+0 isotopologue FE) of label from ^15^N_2_ glutamine in intracellular metabolite (y-axis) pools in each culture condition (x-axis). ns = not significant (unpaired *t*-test for each metabolite). **(D)** Schema depicting derivation of Metabolite Labeling Score. **(E-G)** Volcano plot of Metabolite Labeling Scores in SXO210 preconditioned in HPLM for 120 hours versus Metabolite Labeling Scores derived from ^15^N_2_ glutamine stable isotope tracing assay for 24 hours in the GSCs **(E)** UTSW63, **(F)** TS516, and **(G)** HK157. Horizontal lines represent *p* value of 0.05, vertical lines represent fold-changes of 1.25. Two-tailed *p* values were determined by unpaired *t*-test. **(H-J)** MSEA of differentially labeled metabolites in SXO210 versus **(H)** UTSW63, **(I)** TS516, and **(J)** HK157 GSC lines. *p* values were determined by the quantitative enrichment analysis algorithm. Data are presented as mean ± standard error of the mean.

To compare metabolic activities of heterocellular glioma explants with GSC monocultures, we performed ^15^N_2_-glutamine stable isotope tracing in HPLM in one SXO GBM model and three GBM GSC models: UTSW63, TS516, and HK157. As comparison of label enrichment between SXOs and GSCs may be compromised by differences in proliferation, de novo glutamine synthesis rates, tissue-specific intercellular metabolic cycles, and other factors, we developed a Metabolite Labeling Score to facilitate comparisons across model systems (Figure 3D). The Metabolite Labeling Score uses natural abundance-corrected fractional labeling information, applies quality filters, and then normalizes to the level of label accumulation from glutamine in glutamate. Comparison of metabolite labeling patterns between SXO210 and each of the three GSC lines revealed differences in multiple classes of metabolites, including purine nucleobases, GDP-mannose, and pyrimidine degradation products (Figures 3E-G). Applying Metabolite Set Enrichment Analysis (MSEA) to differentially labeled metabolites between SXO210 and each GSC line, we identified several pathways commonly altered in explants, including purine metabolism and pyrimidine metabolism (Figures 3H-3J). Together, these data indicate that stable isotope tracing in SXOs may enable identification of TME-dependent metabolic phenotypes not readily identified in GSC monocultures.

### Immune-modulatory purine synthesis is a tumor cell-intrinsic feature of glioma metabolism

We observed that glutamine-dependent labeling of various purine metabolites (Figure 4A) differed between explants and GSCs. Compared to SXO210, the GSCs UTSW63, TS516, and HK157 demonstrated high labeling in adenosine and hypoxanthine (Figures 4B and 4C), indicating that degradation of purine nucleotides generated via the de novo purine synthesis pathway is enhanced in tumor cells. Purine nucleotides, such as adenosine monophosphate (AMP) and guanosine monophosphate (GMP), can be synthesized de novo through a multi- enzyme process reliantI e on glutamine as a nitrogen donor.^52^ Purine nucleotides can also be degraded to form nucleosides and nucleobases, including adenosine and hypoxanthine^52^.

**Figure 4.**
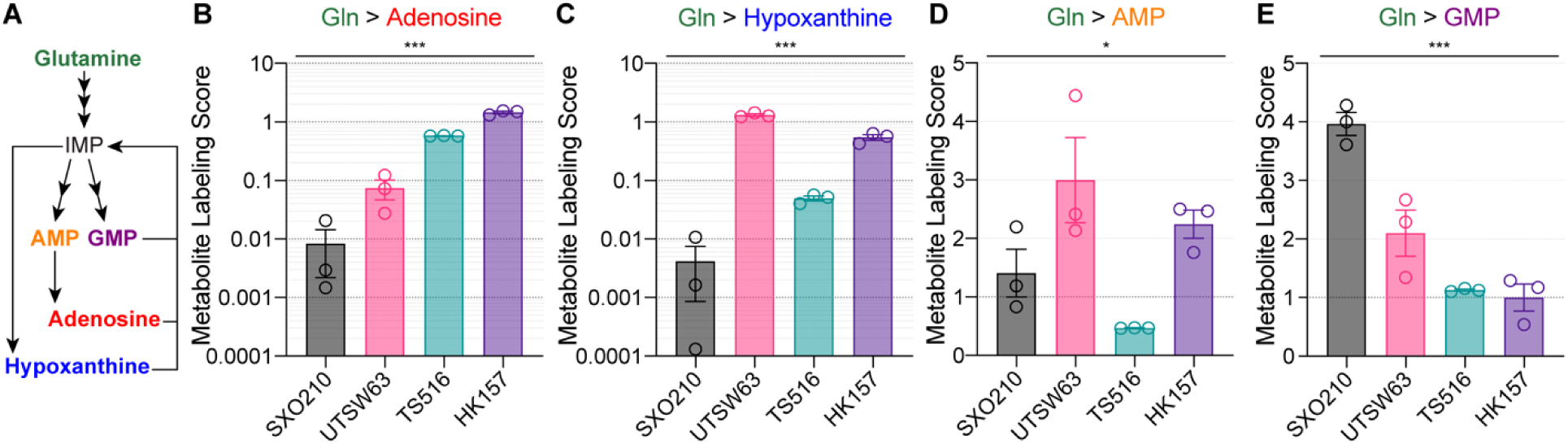
Stable isotope tracing in SXOs captures tumor cell-intrinsic immune-modulatory purinergic metabolism. **(A)** Schematic of glutamine-dependent synthesis of purine nucleotides and their degradation products. IMP = inosine monophosphate. AMP = adenosine monophosphate. GMP = guanosine monophosphate. **(B-E)** ^15^N_2_ glutamine stable isotope tracing assay for 24 hours in SXO210 preconditioned in HPLM for 120 hours and UTSW63, TS516, and HK157 GSC lines. Metabolite Labeling Score in the purine degradation products **(B)** adenosine and **(C)** hypoxanthine and the purine nucleotides **(D)** adenosine monophosphate (AMP) and **(E)** guanosine monophosphate (GMP). *n* = 3 for all groups. \**p* < 0.05, \*\*\**p* < 0.001 (one-way ANOVA). Data are presented as mean ± standard error of the mean.

Extracellular adenosine acts as an important immunosuppressive metabolite through its binding to adenosine receptors on multiple cell types^52–54^. In glioma, accumulation of adenosine through overflow metabolism or extracellular degradation of ATP has been implicated in immune suppression and tumor progression^52,55^. Elevated glutamine-dependent synthesis of adenosine in GSCs could not be explained by higher rates of de novo purine synthesis in these cells, as glutamine labeling of the nucleotide AMP was similar (Figure 4D) and GMP was higher (Figure 4E) in SXOs relative to GSCs. These data indicate that tumor cells make substantial contributions to immunosuppressive adenosine synthesis in the glioma TME. Further, our findings show that comparing metabolic patterns in glioma explants and GSCs can be used to determine glioma cell-intrinsic contributions to brain tumor metabolism.

### Tracing studies in glioma explants reveal stromal production of glycosylation substrates

We also found that glutamine-dependent GDP-mannose synthesis was consistently downregulated in GSCs compared to SXOs (Figures 3E-G and 5A-B). Guanosine diphosphate-mannose (GDP-mannose) is the major mannosyl donor for glycosylphosphatidylinositol anchor synthesis, O-mannosylation, and N-linked glycosylation^56^. Therefore, our data implied that stromal cells in the glioma TME may play an important role in generating metabolic substrates for protein glycosylation. *GMPPA* and *GMPPB* encode GDP-mannose pyrophosphorylase enzymes in humans, which catalyze the conversion of the purine nucleotide guanosine triphosphate and mannose-1-phosphate to GDP-mannose^56,57^ (Figure 5A). GDP-mannose was not detectable in GSCs, while a peak was detectable and quantified in SXO210 (Figure 5C). We analyzed our previously published dataset of RNA sequencing in the three GSC lines^58^ and added analysis of immortalized, nonmalignant astrocytes, NHA. *GMPPA* and *GMPPB* were more highly expressed in NHAs relative to GSCs (Figures 5D and 5E), suggesting that stromal astrocytes drive GDP-mannose synthesis in glioma. Indeed, steady-state levels of GDP-mannose were much higher in NHA astrocytes versus TS516 GSCs (Figure 5F). Both *GMPPA* (Figures 5G and 5H) and *GMPPB* (Figures 5I and 5J) were robustly expressed in SXO210 and normal human brain tissues. Taken together, these data establish that comparative stable isotope tracing in glioma explant and GSC models can be used to identify tumor cell-extrinsic metabolic activities in the glioma TME.

**Figure 5.**
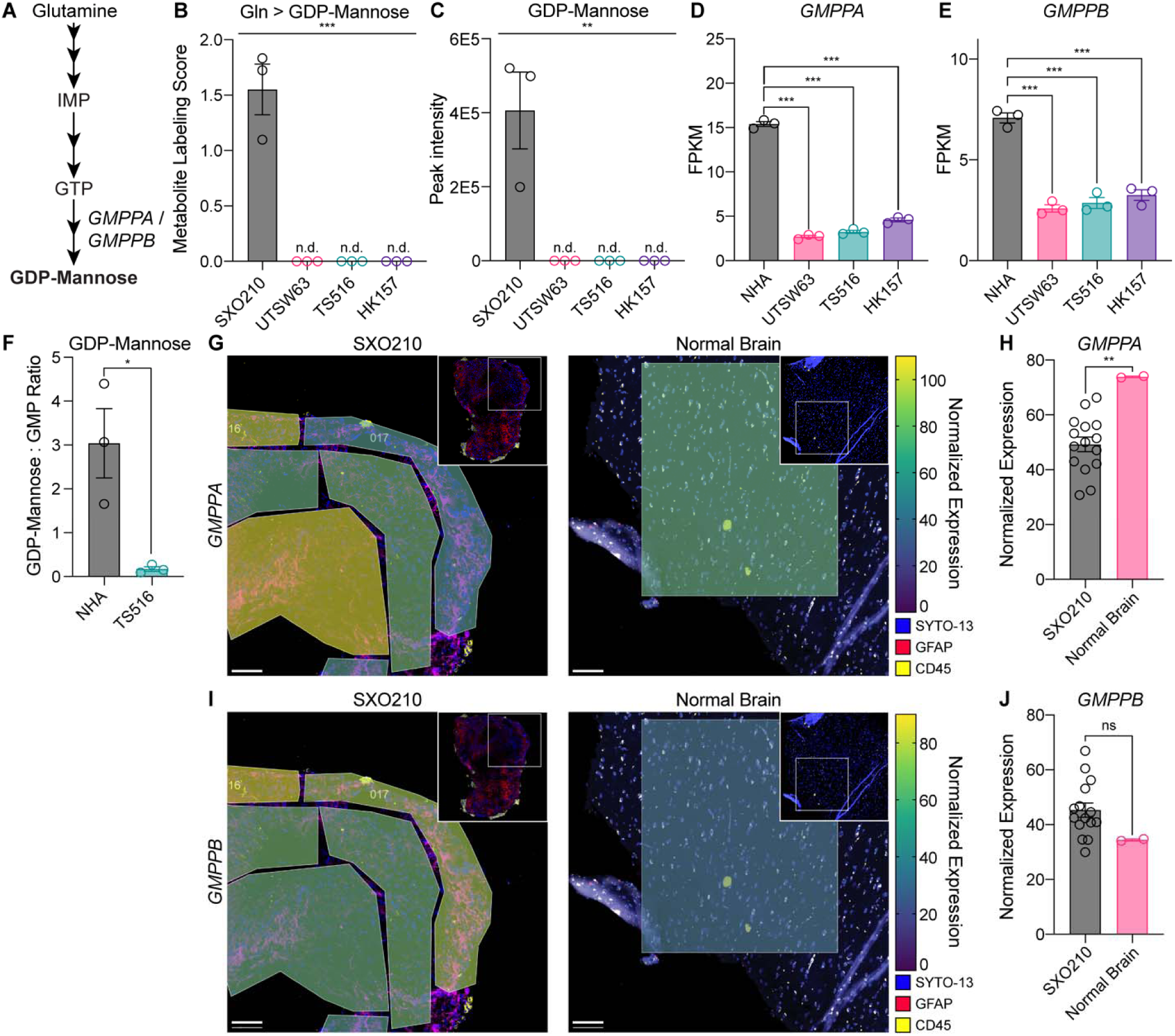
SXO tracing reveals GDP-mannose synthesis by stromal astrocytes in the glioma TME. **(A)** Schematic of de novo purine and guanosine diphosphate-mannose (GDP-mannose) synthesis pathways. IMP = inosine monophosphate. GTP = guanosine triphosphate. **(B)** ^15^N_2_ glutamine stable isotope tracing assay for 24 hours in SXO210 preconditioned in HPLM for 120 hours and UTSW63, TS516, and HK157 GSCs. Metabolite Labeling Score for GDP-mannose. *n* = 3 for all groups. \*\*\**p* < 0.001 (one-way ANOVA). n.d. = not detected. **(C)** Steady-state quantification for GDP-mannose. *n* = 3 for all groups. \*\**p* < 0.01 (one-way ANOVA). n.d. = not detected. **(D-E)** RNA sequencing of NHA immortalized astrocytes and GSC lines, representing fragments per kilobase of transcript per million mapped reads (FPKM) for **(D)** *GMPPA* and **(E)** *GMPPB* genes. *n* = 3 for all groups. ∗∗∗*p* < 0.001 (unpaired *t*-test). **(F)** Amide- ^15^N glutamine stable isotope tracing assay for 18 hours in NHA and TS516 cells. *n* = 3. *p* < 0.05 (unpaired *t*-test). **(G-J)** Heatmap of **(G)** *GMPPA* or **(I)** *GMPPB* expression by geometric mean normalized NanoString sequencing data in representative ROIs of SXO210 explants and normal human brain tissue. Scale bar = 100 μm. Quantification of geometric mean normalized expression in all ROIs of **(H)** *GMPPA* and **(J)** *GMPPB*. Data are presented as mean ± standard error of the mean.

### Pyrimidine degradation reflects the mesenchymal cell state in glioma

Pyrimidine nucleotides represent another class of metabolites that we observed to be differentially labeled by glutamine in SXO and GSC models (Figures 3E-J and 6A). Several lines of evidence led us to hypothesize that this difference may be related to specific glioma cell transcriptional states. GBMs contain cells arising from distinct genetic subclones that display several recurrent transcriptional phenotypes, including a mesenchymal-type associated with loss-of-function mutations in the *NF1* gene^59^. Previous work has linked pyrimidine degradation pathway activity with mesenchymal cell identity in other cancer contexts^60^. Given that the SXO210 model was derived from a GBM with an *NF1* loss-of-function mutation^61^ (Figure 2Z), we hypothesized that SXO210 organoids may be enriched for mesenchymal-type glioma cells that actively degrade pyrimidine nucleotides produced by the glutamine-dependent de novo synthesis pathway. The pyrimidine nucleotide uridine monophosphate (UMP) can be synthesized de novo from glutamine, aspartate, and bicarbonate or salvaged from uridine^52^.

Pyrimidine degradation occurs through multiple steps, the first of which involves conversion of uridine to uracil. Uracil can then be converted to dihydrouracil (DHU) by dihydropyrimidine dehydrogenase, an enzyme encoded by *DPYD*^52^. To test our hypothesis, we first evaluated glutamine-dependent UMP synthesis. SXO210 and GSC cultures exhibited similar glutamine label accumulation in UMP, indicating comparable rates of de novo pyrimidine synthesis (Figure 6B). In contrast, glutamine labeling of pyrimidines downstream of UMP displayed substantial heterogeneity across models. Label from glutamine variably accumulated in UDP-hexose (Figure 6C), while all models exhibited label accumulation in the pyrimidine salvage substrate uridine (Figure 6D). While label from glutamine was present in uridine across all lines, only SXO210 exhibited label accumulation in uracil (Figure 6E). This result indicated that only the SXO210 model displayed robust pyrimidine degradation. Consistent with the idea that this metabolic activity may be driven by mesenchymal glioma cells enriched in SXO210 explants, the UTSW63, TS516, and HK157 GSC lines expressed low or undetectable levels of the mesenchymal marker CD44 (Figure 6F).

**Figure 6.**
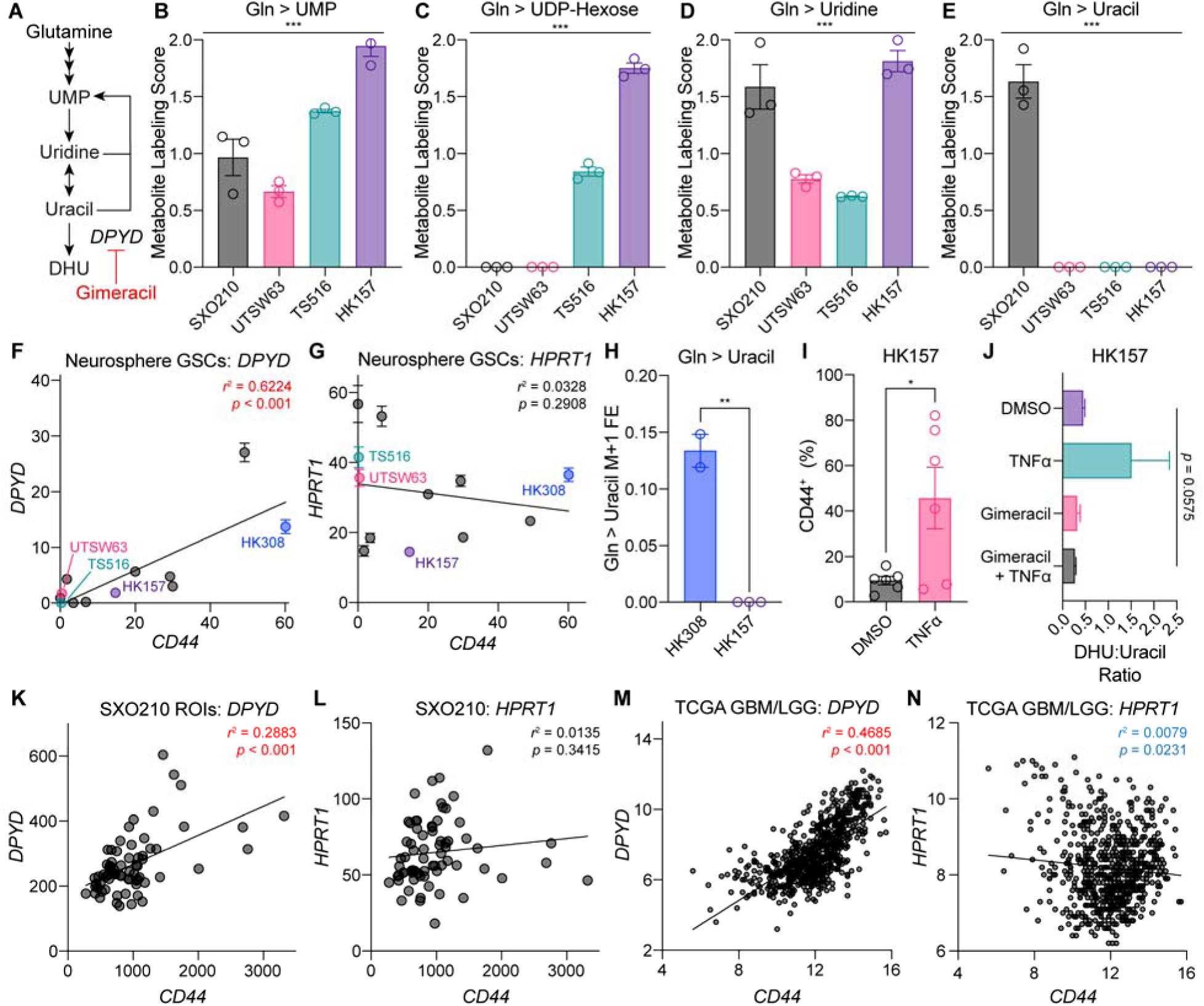
Metabolic features defining the mesenchymal transcriptional subtype of glioma in SXOs. **(A)** Schematic of pyrimidine synthesis and degradation pathways. UMP = uridine monophosphate. DHU = dihydrouracil. **(B-E)** ^15^N_2_ glutamine stable isotope tracing assay for 24 hours in SXO210 preconditioned in HPLM for 120 hours and UTSW63, TS516, and HK157 GSC lines. Metabolite Labeling Score for **(B)** UMP, **(C)** UDP-hexose, **(D)** uridine, **(E)** and uracil. *n* = 3 for all groups. \*\*\**p* < 0.001 (one-way ANOVA). **(F-G)** Correlation between expression of *CD44* and **(F)** *DPYD* or **(G)** *HPRT1* in the GSC lines BT054, HK157, HK211, HK252, HK308, MGG152, TS516, TS603, UTSW5, UTSW63, and UTSW71 by RNA sequencing. Expression values are shown in Fragments Per Kilobase of transcript per Million mapped reads (FPKM). *r* = Pearson’s correlation coefficient. *n* = 3 per line. **(H)** Amide-^15^N glutamine stable isotope tracing assay for 18 hours in HK308 and HK157. *n* = 2 in HK308, *n* = 3 in HK157. \*\**p* < 0.01 (unpaired *t*-test). FE = fractional enrichment. **(I)** Flow cytometry quantification of CD44 in HK157 cells treated for 7 days with DMSO or TNFα. ∗p < 0.05 (unpaired *t*-test). **(J)** Ratio of DHU to uracil in HK157 cells treated for 7 days with DMSO, 10 ng/mL TNFα, 30 µM gimeracil, or 10 ng/mL TNFα and 30µM gimeracil. *p* value was determined by one-way ANOVA**. (K-L)** Correlation between geometric mean normalized expression of *CD44* and **(K)** *DPYD* or **(L)** *HPRT1* in SXO210 ROIs by NanoString sequencing. *p* values were determined by simple linear regression. **(M-N)** Correlation between expression of *CD44* and **(M)** *DPYD* or **(N)** *HPRT1* in The Cancer Genome Atlas (TCGA) GBM and Lower-Grade Glioma (LGG) datasets. The results shown here are based upon data generated by the TCGA Research Network. Expression values are log_2_ transformed and normalized by RSEM. *p* value was determined by simple linear regression. Data are presented as mean ± standard error of the mean.

To investigate whether pyrimidine degradation is associated with the mesenchymal subtype of GBM, we leveraged an RNA sequencing dataset of GSC lines^58^. We found a strong correlation between expression of the mesenchymal marker *CD44* and *DPYD*, but not with a cognate enzyme in purine nucleotide degradation, *HPRT1* (Figures 6F and 6G). Using the GSC line with the highest level of CD44 expression in this dataset (HK308) and CD44-low HK157 cells, we performed amide-^15^N glutamine stable isotope tracing and confirmed that glutamine label accumulated in uracil in mesenchymal HK308 cells but not non-mesenchymal HK157 cells (Figure 6H). Tumor necrosis factor-alpha (TNFα) is an inflammatory cytokine that can induce epithelial-to-mesenchymal transition^62^. Treating HK157 cells with TNFα induced mesenchymal identity, as marked by CD44 upregulation (Figure 6I). Treating HK157 cells with TNFα also increased the ratio of DHU:uracil, a metabolic marker of pyrimidine degradation^60^ (Figure 6J), which was reversed by the DPYD inhibitor gimeracil (Figure 6J). These findings indicate that mesenchymal differentiation of GSCs is sufficient to activate pyrimidine degradation.

Intratumoral heterogeneity of transcriptional phenotypes is a common feature of GBMs^59^, so we examined the association of the mesenchymal phenotype with *DPYD* expression in individual ROIs of SXO210 explants via spatial transcriptomics. Regions with higher CD44 expression were associated with higher expression of *DPYD* (Figure 6K), but not *HPRT1* (Figure 6L) expression. To test whether this association was broadly relevant in glioma, we turned to The Cancer Genome Atlas studies of lower-grade glioma (LGG)^44^ and GBM^45^. Consistent with our cell culture and SXO studies, *CD44* expression by RNA sequencing in patient samples was positively correlated with expression of *DPYD* (Figure 6M) but not *HPRT1* (Figure 6N). Taken together, our data establish that metabolic hallmarks of GBM transcriptional subtypes are reflected in stable isotope tracing assays of glioma explants.

## DISCUSSION

Recent advances in mass spectrometry technology and metabolite probe development have produced deeper understanding of tumor metabolism. In glioma, the discovery of recurrent hotspot mutations in *IDH1* and *IDH2* genes^63,64^ has provided added rationale to investigate the association between altered metabolism and oncogenesis. Investigation of tumor-intrinsic metabolism in glioma has yielded insights into therapeutic vulnerabilities, including de novo pyrimidine synthesis^3,4,7,17^, purine synthesis^8,65^, glutathione metabolism^9,10^, dopaminergic signaling^11^, malate dehydrogenase activity^12^, and threonine metabolism^13^. Glioma cells further have been shown to alter cell-intrinsic nucleotide metabolism to mediate chemo- and radio- resistance^14–16,65,66^. Additional research has revealed critical contributions of intercellular crosstalk in the TME to glioma biology and metabolism. For example, altered metabolism of glioma cells has been linked to epigenetic rewiring that suppresses antitumor immunity^67,68^ and TME lymphatic endothelial-like cells have been shown to regulate glioma cell cholesterol metabolism^69^.

Steady-state metabolomics analyses of primary tissue samples have provided key insights into TME-dependent contributions to glioma metabolism^70–73^. These studies measure total metabolite levels but are limited in their ability to assess metabolic pathway activity^74^. In comparison, stable isotope tracing techniques offer the ability to monitor dynamic metabolic processes. Intraoperative stable isotope infusions in patients with gliomas have been reported using ^13^C_6_-glucose or hyperpolarized ^13^C-labeled metabolite probes^20–23^. These techniques, however, are expensive, may require perioperative intervention, involve intricate coordination between clinicians and researchers, and typically investigate a single metabolite tracer per patient.

Studies of metabolism involving banked tissue specimens or tracer infusions in patients are complemented by cell culture and murine disease models. Employing such models obviates the need for clinical coordination and creates opportunities to apply multiplexed or high-throughput approaches to assess tumor metabolism. However, conventional cell culture models often lack the metabolic and cellular complexity of the tumor microenvironment (TME), and common incorporation of non-physiologic nutrient conditions limits translational relevance. While orthotopic murine models address some of these limitations, they, too, are technically challenging and expensive—particularly for isotope tracing—and are constrained by the use of immunocompromised hosts, precluding study of immune-metabolic crosstalk^75,76^. These challenges underscore the need for tractable, physiologically faithful ex vivo models that recapitulate the metabolic and immunologic context of human gliomas^31^.

We present a method that integrates defined, physiologic culture conditions with stable isotope tracing to interrogate glioma metabolism. A key feature of our approach is the use of an adapted formulation of HPLM^27^, a medium that approximates the nutrient composition of human plasma. Importantly, we show that this SXO-adapted HPLM formulation supports glioma explant culture while preserving native features of tumor architecture and cell composition. When combined with spatial transcriptomics and isotope tracing, this approach captures metabolic programs that reflect both tumor-intrinsic and microenvironmental activity. Importantly, glioma explants cultured in HPLM exhibit immune and metabolic transcriptional responses not observed in traditional media, suggesting that nutrient context actively shapes the tumor transcriptional state. These responses support the application of HPLM for the discovery of metabolic programs that govern interactions between tumor cells and the TME—features that are lost in neurosphere GSC monocultures.

While GSC models provide valuable insights into tumor-intrinsic metabolic programs, surveying metabolism in glioma explants allows these same programs to be examined in the context of immune and stromal interactions. Our tissue explant model reveals stromal contributions to glioma metabolism that are not evaluable in GSC monocultures. Indeed, metabolites that are regulated by stromal cell metabolism, such as astrocyte-derived GDP-mannose, can be effectively studied using our comparative stable isotope tracing approach. This approach has the advantage of directly measuring both metabolite levels and metabolic activities, rather than relying on expression of metabolic enzymes as a proxy. This is particularly relevant to the case of GDP-mannose production, as both our study and another^77^ report expression of constituent enzymes of its synthetic pathway in GBM cells.

The heterogeneity of GBM and underlying transcriptional subtypes represents an important aspect of the biology of these tumors^59^. Our approach captures metabolic features of GBM transcriptional subtypes, including increased flux through the pyrimidine degradation pathway in mesenchymal glioma cells. Building on prior work in other cancers^60^, we demonstrate that inducing mesenchymal cell state transitions in GSCs is sufficient to activate pyrimidine degradation. Our study therefore deepens our understanding of connections between metabolic plasticity and inflammatory and transcriptional cues within the glioma microenvironment.

By enabling analysis of both cell-autonomous and non-cell-autonomous metabolic programs, our platform enhances the resolution with which glioma metabolism can be interrogated. The ability to maintain intercellular interactions in a nutrient context that is physiologically accurate positions this approach to uncover actionable metabolic dependencies that are tightly coupled to the glioma TME. Stable isotope tracing in HPLM provides a functional platform for evaluating metabolic phenotypes that may not emerge in non-physiologic conditions, opening avenues for therapeutic discovery based on nutrient-sensitive vulnerabilities.

Our study marries advances in tissue culture, stable isotope tracing, and glioma organoid modeling. We acknowledge that this work is subject to several limitations. First, although numerous explants were used in the SXO stable isotope tracing analysis, they originated from a single patient. Comparison of SXOs to GSCs was performed using GSC lines from different parental tumors, limiting explanatory power potentially gained from evaluating heterocellular SXOs and GSC monocultures derived from a common tumor. Future studies will focus on establishing SXOs and GSCs from the same parental tumor to improve discrimination of TME-dependent features of glioma metabolism. We also performed stable isotope tracing at a single timepoint, thereby not allowing for formal evaluation of flux in these models. Tracing with only ^15^N_2_ glutamine, additionally, may not reveal important metabolic differences that could be apparent with tracing by other carbon or nitrogen metabolic substrates^78^. Additional studies investigating the metabolism of other substrates could provide deeper insights into metabolic activities specific to the glioma TME that affect tumor cell viability and growth. Despite these limitations, our findings lay the groundwork for improving the physiological relevance of in vitro studies of glioma metabolism.

## REQUIRED STATEMENTS

### Ethics

All patient specimens were collected following ethical and technical guidelines on the use of human samples for biomedical research at University of Texas Southwestern Medical Center under protocols approved by the University of Texas Southwestern Medical Center’s IRB (STU 022011-070 and STU 092014-026). Informed consent was obtained from all individual participants included in the study.

## Funding

This project was supported by National Institutes of Health (NIH) grants R01CA258586 and R01CA289260 to S.K.M. and K.G.A., P50CA165962 and U19CA264504 to S.K.M., and F30CA271634 to M.R.S. This work was also supported by an award from Oligo Nation to S.K.M. and K.G.A. and by Cancer Prevention and Research Institute of Texas (CPRIT) grants RR190034, RP230344, RP2400489, RP250278 to S.K.M. L.G.Z., T.P.M., and the Children’s Research Institute Metabolomics facility are supported by the Cancer Prevention Research Institute of Texas (CPRIT Core Facilities Support Award RP240494). Y.X. was supported by an NIH/NCI grant (K99CA277576) and a Human Frontier Science Program postdoctoral fellowship award (LT0018/2022-L).

### Authorship

Conceptualization: MRS, MES, SKM, KGA; Data curation: MRS, MES, LCG, BCS, WHH, SSO; Formal Analysis: MRS, MES, LCG, BCS, WHH, SSO, TER; Funding acquisition: MRS, SKM, KGA; Investigation: MRS, MES, LCG, BCS, WHH, SSO, LGZ, MSMS, EYM, POZ, BCL; Methodology: MRS, MES, LCG, BCS, WHH, SSO, LGZ, MSMS, EYM, POZ, BCL, TPM; Resources: POZ, BCL, KGA; Software: MRS; Supervision: JNR, TPM, RJD, SKM, KGA; Validation: MRS, MES, TER; Visualization: MRS, MES, LCG, WHH; Writing – original draft: MRS, MES; Writing – review & editing: MRS, YX, DDS, JNR, SKM, KGA

### Data Availability

Raw histopathology images and NanoString morphology scans will be shared by the lead contact upon request. Metabolomics datasets will be made available on the NIH National Metabolomics Data Repository. NanoString sequencing results will be made available in the NCBI Gene Expression Omnibus. RNA sequencing of GSC lines was previously published by Wu et al.^58^, but will be included along with RNA sequencing analysis of NHA for ease of reanalysis in the NCBI Gene Expression Omnibus. Any additional information required to reanalyze the data reported in this paper is available from the lead contact upon request.

