## Supplementary Table 1 for "Stable isotope tracing in human plasma-like medium reveals metabolic and immune modulation of the glioblastoma microenvironment"

| **Sample ID** | **Model** | **Age** | **Sex** | **Brain Region** | **WHO Grade** | **Diagnosis** | **Primary/**  **Recurrent** |
| --- | --- | --- | --- | --- | --- | --- | --- |
| UTSW63 | Cell Line | 71 | M | Right Temporal | 4 | GBM, IDH-wildtype | Primary |
| SXO210 | SXO | 68 | M | Right Temporal | 4 | GBM, IDH-wildtype | Primary |
| Normal Brain | Primary Tissue | 42 | F | Right Temporal | N/A | Intractable epilepsy | N/A |

**Supplementary Table 1.** Clinical characteristics of primary tumor or normal brain specimens used for SXO and GSC lines derived in this study. “Brain Region” corresponds to the location of the tumor or brain at resection. “Diagnosis” corresponds to the results of histopathologic features and molecular studies rendered by a clinical pathologist at resection. Abbreviations: M, male; F, female, GBM, glioblastoma; IDH, isocitrate dehydrogenase; SXO, surgically explanted organoid. IDH status of each primary tumor sample was assessed via immunohistochemistry, next-generation DNA sequencing, or both.
